# Leptin-mediated suppression of food intake by conserved *Glp1r*-expressing neurons prevents obesity

**DOI:** 10.1101/2021.12.10.472115

**Authors:** Alan C. Rupp, Abigail J. Tomlinson, Alison H. Affinati, Cadence True, Sarah R. Lindsley, Melissa A. Kirigiti, Alexander MacKenzie, Chien Li, Lotte Bjerre Knudsen, David P. Olson, Paul Kievit, Martin G. Myers

**Affiliations:** Department of Internal Medicine, University of Michigan, Ann Arbor, MI, USA; Oregon National Primate Research Center, Beaverton, OR, USA; Novo Nordisk Research Center, Seattle, WA, USA; Department of Pediatrics, University of Michigan, Ann Arbor, MI

## Abstract

The adipose-derived hormone leptin acts via its receptor (LepRb) in the brain to control energy balance. A previously unidentified population of GABAergic hypothalamic LepRb neurons plays key roles in the restraint of food intake and body weight by leptin. To identify markers for candidate populations of LepRb neurons in an unbiased manner, we performed single-nucleus RNA-sequencing of enriched mouse hypothalamic LepRb cells, as well as with total hypothalamic cells from multiple mammalian species. In addition to identifying known LepRb neuron types, this analysis identified several previously unrecognized populations of hypothalamic LepRb neurons. Many of these populations display strong conservation across species, including GABAergic *Glp1r*-expressing LepRb (LepRb^Glp1r^) neurons that express more *Lepr* and respond more robustly to exogenous leptin than other LepRb populations. Ablating LepRb from these cells provoked hyperphagic obesity without impairing energy expenditure. Conversely, reactivating LepRb in *Glp1r*-expressing cells decreased food intake and body weight in otherwise LepRb-null mice. Furthermore, LepRb reactivation in GABA neurons improved energy balance in LepRb-null mice, and this effect required the expression of LepRb in GABAergic *Glp1r*-expressing neurons. Thus, the conserved GABAergic LepRb^Glp1r^ neuron population plays crucial roles in the control of food intake and body weight by leptin.

## Introduction

The ongoing obesity pandemic represents an enormous challenge to human health and longevity (1). Identifying therapeutic targets to combat obesity will require understanding the physiologic systems that modulate feeding and maintain appropriate body weight. The adipose-derived hormone, leptin, controls body weight by signaling the repletion of body fat stores to hypothalamic neurons that express the long isoform of the leptin receptor (LepRb) (2, 3). Insufficient leptin action, as following the reduction of fat stores by prolonged caloric restriction, increases hunger and inhibits energy-intensive neuroendocrine systems (4, 5). Similarly, humans and animals lacking leptin or LepRb are unable to sense adipose energy stores and thus exhibit voracious feeding and decreased energy expenditure despite their severe obesity.

Hypothalamic LepRb neurons that control energy balance represent important potential targets for obesity therapy, but the cell types most important for the control of body weight by direct leptin action remain to be defined. Although arcuate nucleus (ARC) proopiomelanocortin (*Pomc*)-expressing and distinct agouti-related peptide (*Agrp*)-expressing neurons play important roles in energy balance (6, 7), the early developmental ablation of *Lepr* from these cells only modestly impacts feeding and body weight (8-10). While developmental compensation may partially conceal the role for LepRb in AgRP neurons following its early ablation (9, 11, 12), animals lacking LepRb throughout the body or in the hypothalamus from early developmental times exhibit dramatic hyperphagic obesity despite any compensation for their lack of LepRb in AgRP neurons (13). Thus, other (non-AgRP) hypothalamic LepRb neurons not subject to the same level of developmental compensation as AgRP neurons must play major roles in leptin action.

Several lines of evidence suggest the importance of an as-yet unidentified set of GABAergic LepRb neurons for the control of food intake and body weight; these likely reside in the dorsomedial hypothalamic nucleus (DMH) (10, 14-16). To identify potentially important hypothalamic LepRb neuron populations, we utilized single-nucleus RNA-sequencing (snRNA-seq) to define hypothalamic LepRb neuron populations in an unbiased manner. In addition to identifying previously-described types of LepRb neurons, this analysis revealed several previously unknown LepRb populations, including those marked by *Glp1r, Tbx19*, and *Foxb1*. Of these, we studied a conserved GABAergic *Glp1r*-expressing population of LepRb neurons that reside mainly in the DMH. These LepRb^Glp1r^ neurons play major roles in the leptin-mediated control of food intake and body weight.

## Results

To identify LepRb-expressing hypothalamic cell populations in an unbiased way, we employed the LepRb-specific *Lepr*^*Cre*^ allele in combination with a reporter allele that Cre-dependently expresses a nuclear-localized Sun1-sfGFP fusion protein in LepRb cells (LepRb^Sun1-sfGFP^ mice, Fig. 1A, S1). We FACS-sorted GFP-containing nuclei from LepRb^Sun1-sfGFP^ hypothalamic tissue and subjected the nuclei to snRNA-seq using the 10x Genomics 3’ system (Fig. 1B). After removing contaminants (including neurons derived from neighboring non-hypothalamic brain areas (Fig. S2)), we used graph-based clustering of the remaining 2879 nuclei to identify 18 populations of hypothalamic LepRb neurons (LepRb-Sun1 populations) (Fig. 1C). Most of these populations exhibited dozens of unique marker genes (Fig. 1D). A small number of LepRb-Sun1 populations (e.g., GABA, GLU1, GLU2, and GLU3) displayed no clear markers and weak transcriptional enrichment, however; these likely represent collections of GFP-negative contaminants and/or amalgamations of smaller populations of LepRb neurons.

**Fig. 1.**
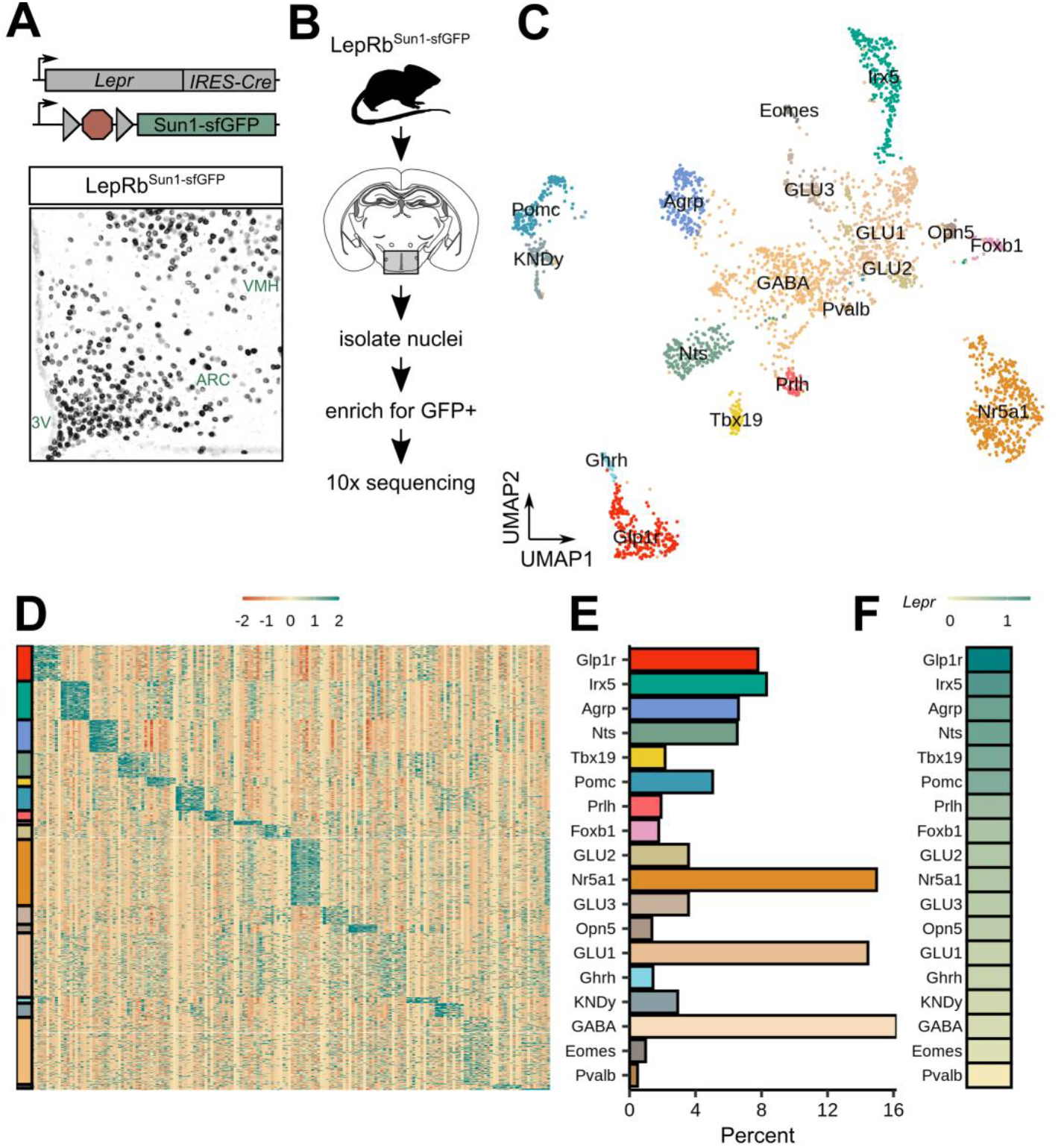
snRNA-seq of FACS-enriched hypothalamic LepRb neurons defines known and novel LepRb neuron populations. (**A**) Genetic diagram for the LepRb^Sun1-sfGFP^ mouse line (top), and representative image showing GFP-immunoreactivity (-IR) (black) in the mediobasal hypothalamus of a LepRb^Sun1-sfGFP^ mouse (bottom) 3V, third cerebral ventricle. (**B**) Experimental diagram for isolating LepRb nuclei from the hypothalamus for snRNA-seq. (**C**) UMAP projection of all 2879 hypothalamic LepRb^Sun1-sfGFP^ neuronal nuclei, colored by cluster. (**D**) Scaled expression of top marker genes across all cells; colors on left correspond to colors of populations as in C, E. (**E**) Percent of cells that map to each cluster. (**F**) Mean *Lepr* expression, normalized to the average expression in glia.

These 18 snRNA-seq-defined LepRb-Sun1 populations included previouslydescribed discrete hypothalamic LepRb cell types, validating our snRNA-seq approach to identifying populations of hypothalamic LepRb neurons. These previously-described populations included those marked by the expression of *Ghrh* (LepRb^Ghrh^ cells, which reside in the ARC); *Nr5a1* (LepRb^Nr5a1^; ventromedial hypothalamic nucleus (VMH)); *Irx5* (LepRb^Irx5^ neurons; ventral premammillary nucleus (PMv)); *Agrp* (LepRb^AgRP^ neurons; ARC); *Prlh* (LepRb^Prlh^ neurons; DMH); *Nts* (LepRb^Nts^ neurons; lateral hypothalamic area (LHA) and DMH); *Pomc* (LepRb^Pomc^ neurons; ARC); and *Kiss1/Tac2/Pdyn* (LepRb^KNDy^ neurons; ARC) (10, 17-23). For all LepRb populations, the marker genes denoted represent those expressed across many neurons of the designated population and display expression that is largely restricted to this population; some populations express additional strong marker genes (e.g., *Npy* in LepRb^Agrp^ cells).

We also identified six previously unstudied populations of hypothalamic LepRb neurons with robust gene expression markers, including those marked by the expression of *Glp1r, Tbx19, Foxb1, Opn5, Eomes*, and *Pvalb* (designated LepRb^Glp1r^, LepRb^Tbx19^, LepRb^Foxb1^, LepRb^Opn5^, LepRb^Eomes^, and LepRb^Pvalb^ neurons, respectively) all of which exhibited robust gene expression markers. Notably, most LepRb-Sun1 populations expressed substantially more *Lepr* than glial cells (which do not express LepRb ((24, 25) and S2F)). LepRb^Glp1r^ neurons exhibited higher *Lepr* expression than any other LepRb population (Fig. 1E).

To learn more about the potential regulation of LepRb-Sun1 neuron populations by leptin, we used translating ribosome affinity purification with RNA-sequencing (TRAP-seq) data from LepRb^EGFP-L10a^ mice, which express a GFP-tagged L10a ribosomal subunit in LepRb neurons (Fig. 2A)(26). We found that many LepRb TRAP-seq-enriched transcripts mapped strongly to specific LepRb-Sun1 populations (Fig. 2B), suggesting that the gene expression profiles of LepRb-Sun1 populations could be used to reveal cell type-specific gene expression programs. Indeed, many leptin-regulated transcripts identified by LepRb neuron-specific TRAP-seq (Fig. 2C) displayed variable expression across the LepRb-Sun1 populations (Fig. 2D). Hence, we generated an aggregate score for the enrichment of leptin-regulated transcripts for each LepRb-Sun1 population (Fig. 2E), which suggested that transcriptional responses to exogenous leptin are biased toward LepRb^Glp1r^, LepRb^AgRP^, LepRb^Pomc^, LepRb^Ghrh^, and LepRb^Tbx19^ neuron populations.

**Fig. 2.**
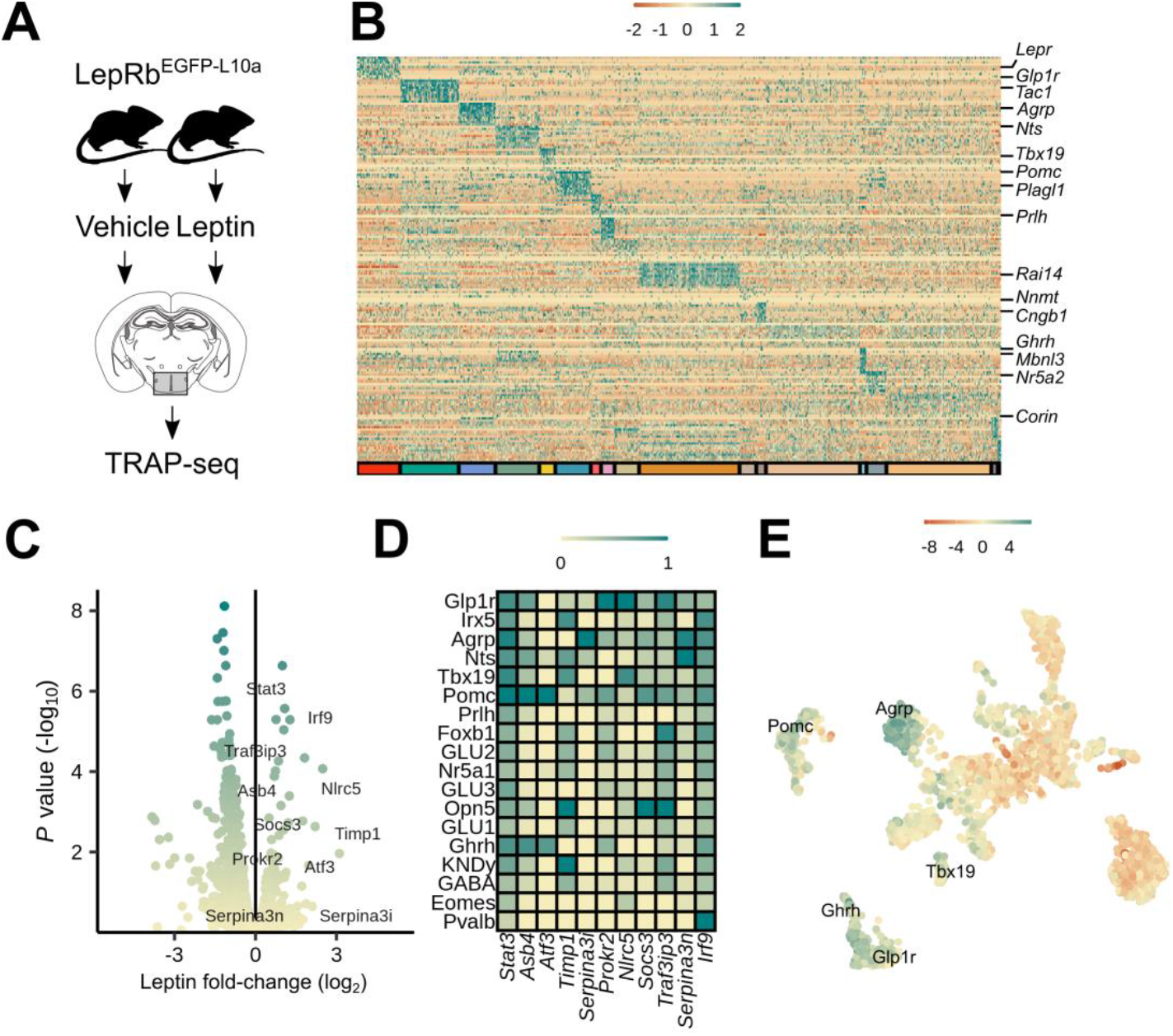
Biased mapping of leptin-regulated transcripts toward specific LepRb-Sun1 populations. (**A**) Experimental diagram for published LepRb TRAP-seq (data available at GSE162603 (42)). (**B**) Relative expression of TRAP-seq enriched genes in specific LepRb-Sun1 populations. (**C**) Volcano plot of leptin-regulated changes in gene expression in TRAP-seq analysis. (**D**) Average expression of representative highly leptin-regulated genes across all LepRb-Sun1 clusters. (**E**) UMAP projection of all LepRb-Sun1 cells, colored by loading of leptin-regulated genes.

To determine whether these LepRb-Sun1 populations correspond to specific mediobasal hypothalamic cell types and whether these cell types are conserved in other species, we mapped neurons from our LepRb-Sun1 analysis onto new snRNA-seq-derived cell atlases of the mouse and rat mediobasal hypothalamus that we generated for this purpose, and from our previously-reported macaque mediobasal hypothalamus snRNA-seq dataset (Fig. 3A) (27). These datasets recovered all major cell types in all species in similar proportions (Fig. S3). *Lepr* transcripts in these mediobasal hypothalamic atlases were almost exclusively detected in neurons (Fig. S3N), as in our LepRb-Sun1 dataset (Fig. S2F). Subclustering of the neurons from each species revealed dozens of populations with clearly enriched marker genes (Fig. 3B–D).

**Fig. 3:**
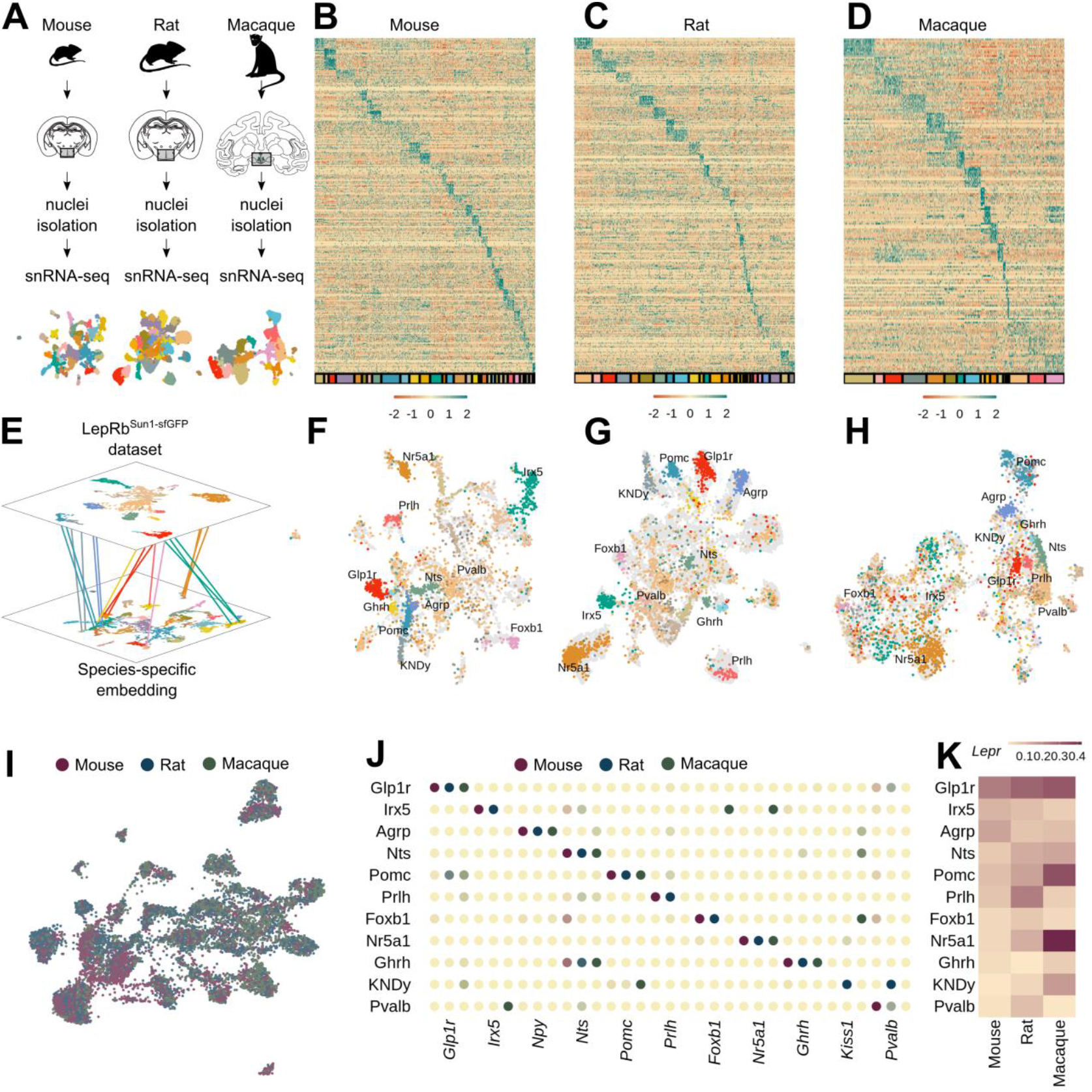
Conservation of LepRb populations in rodents and macaques. (**A**) Diagram for snRNA-seq of mouse, rat, and macaque and UMAP projection of all neurons, colored by cluster. (**B**–**D**) Scaled expression of top marker genes for each species across all cells, colored by population (as in UMAP plots) on the bottom. (**E**) Strategy for identifying orthologous cells across datasets by embedding LepRb-Sun1 cells in UMAP space for hypothalamic cells from each species. (**F**–**H**) UMAP embedding of each species (gray) overlayed with UMAP embedding of LepRb-Sun1 cells (colors and labels are from LepRb-Sun1 populations as in Figure 1; labels are shown for populations that retain strong clustering across species). (**I**) UMAP plot of co-clustered cells from all species, using conserved orthologous genes; species of derivation of each cell is denoted by the color of the cell. (**J**) Expression of marker genes for each conserved cluster across species (colors represent scaled expression ranging from 0–1 for each species). (**K**) Mean *Lepr*/*LEPR* expression, normalized to the average expression in glia.

To identify common hypothalamic LepRb neuron populations across species, we first projected our LepRb-Sun1 cells into the UMAP embeddings of each dataset (Fig. 3E). Many LepRb-Sun1 populations (including LepRb^Glp1r^, LepRb^Agrp^, and LepRb^Pomc^, among others) retained dense associations in all embeddings, suggesting they have clear orthologs in all datasets (Fig. 3F–H). We also co-clustered neurons from the LepRb-Sun1 snRNA-seq with mediobasal hypothalamic neurons from each species, using genes orthologous across species (Fig. 3I, S4). This consensus clustering revealed eleven well-conserved LepRb-Sun1-containing cell clusters that presumably represent *bona fide* LepRb populations in each species. Major marker genes for most of these conserved LepRb populations were maintained across species, albeit with a few exceptions (Fig. 3J).

Interestingly, we observed some population-specific differences in *Lepr* expression by species, including relatively higher *Lepr* expression in LepRb^Agrp^ and LepRb^Irx5^ populations in the mouse than other species, and relatively higher *Lepr* expression in the LepRb^Nr5a1^ and LepRb^Pomc^ populations in the macaque than in rodents (Fig. 3K). Cells from the LepRb^Glp1r^ population exhibited consistently high *Lepr* expression across species, however (Fig. 3K).

Combined, the findings of high *Lepr* expression, robust enrichment of leptin-regulated genes, and conservation across species suggest potentially important roles for the LepRb^Glp1r^ population in leptin action. We thus chose to determine the roles for this LepRb population in leptin action *in vivo*. As expected, the vast majority of *Glp1r/Lepr*-coexpressing cells from our LepRb-Sun1 analysis mapped to the LepRb^Glp1r^ population (Fig. 4A), suggesting that the analysis of *Glp1r*-expressing LepRb neurons (LepRb^Glp1r^ cells) should reveal the function of this population of cells.

**Fig. 4.**
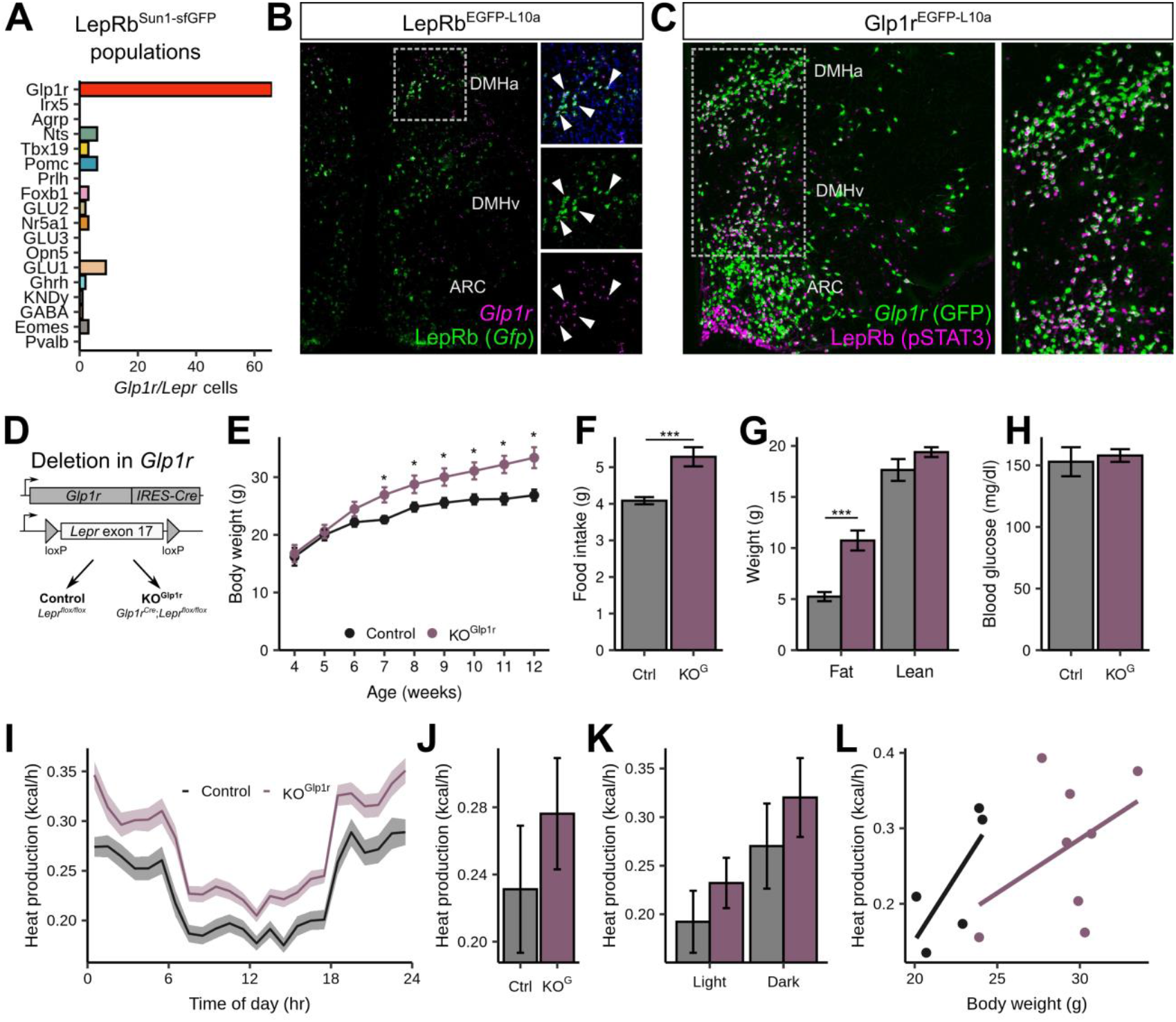
Requirement for *Lepr* in LepRb^Glp1r^ cells for the normal control of food intake and body weight by leptin. (**A**) The number of *Glp1r*-expressing cells across all LepRb^Sun1-sfGFP^ populations. (**B**) Representative image showing *in situ* hybridization for *Gfp* (LepRb, green) and *Glp1r* (magenta) in the hypothalamus of a LepRb^EGFP-L10a^ mouse. (**C**) Representative image showing GFP-IR (green) and pSTAT3-IR (LepRb/pSTAT3, magenta) in the hypothalamus of a leptin-treated Glp1r^EGFP-L10a^ mouse. Inset shows a digital zoom of the boxed region in the larger panel. (**D**–**H**) Generation and analysis of KO^Glp1r^ mice. (**D**) Experimental diagram for generating KO^Glp1r^ and Control animals. (**E**) Body weight, (**F**) food intake, (**G**) body composition, and (**H**) blood glucose in Control (black, *n* = 6–7) and KO^Glp1r^ (purple, *n* = 11) male mice. (**I**–**N**) Measurement of heat production in KO^Glp1r^ mice (*n* = 8) and Control (*n* = 5) male mice. (**I**) Hourly average, (**J**) daily average, (**K**) daily average within light/dark cycle, and (**L**) correlation of heat production and body weight. Statistical significance determined by permutation test. *p<0.05; ***p<0.001.

*In situ* hybridization for *Glp1r* and *Gfp* in LepRb^EGFP-L10a^ mice revealed a substantial population of LepRb^Glp1r^ cells in the DMH (Fig. 4B). Using *Glp1r*^*Cre*^ on a reporter strain that cre-inducibly expresses an EGFP-L10a fusion protein (Glp1r^EGFP-L10a^ mice) similarly revealed the colocalization of leptin-stimulated phosphorylated STAT3-immunoreactivity (pSTAT3-IR, a marker for cell-autonomous leptin action) with GFP in the DMH, with smaller populations of cells in the adjacent ARC and LHA (Fig. 4C, Fig. S5A). We also directly examined the potential colocalization of POMC-IR with pSTAT3-IR and EGFP in the ARC of leptin-treated Glp1r^EGFP-L10a^ mice to rule out potentially confounding effects on POMC neurons during the *Glp1r*^*Cre*^-dependent ablation of *Lepr* in LepRb^Glp1r^ cells (despite the minimal impact of *Lepr* ablation from *Pomc* cells on energy balance (8-10)). As previously reported by others (28), we observed very few POMC cells containing both *Glp1r*^*Cre*^ and pSTAT3 (Fig. S5B). Thus, LepRb^Glp1r^ neurons reside predominantly within the DMH, although this population contains a few (non-POMC) neurons in the ARC and LHA.

To determine roles for LepRb^Glp1r^ cells in leptin action, we crossed *Glp1r*^*Cre*^ onto the *Lepr*^*flox*^ background to generate *Glp1r*^*Cre*^*;Lepr*^*flox/flox*^ (KO^Glp1r^) mice lacking *Lepr* specifically in *Glp1r*-expressing neurons (Fig. 4D). Both male (Fig. 4E–H) and female (Fig. S6A–D) KO^Glp1r^ mice displayed hyperphagic obesity with unperturbed blood glucose. Interestingly, while LepRb-null mice exhibit severely reduced energy expenditure (29), KO^Glp1r^ mice displayed a trend toward increased energy expenditure compared to lean controls as would be expected with increased body weight (Fig. 4I–L, S6–7), suggesting that leptin action via LepRb^Glp1r^ cells primarily controls food intake (rather than energy expenditure).

We also examined the effect of restoring *Lepr* expression in *Glp1r* cells on an otherwise *Lepr*-null background in *Glp1r*^*Cre*^*;Lepr*^*LSL*^ (Fig. 5A, Re^Glp1r^) mice. This maneuver substantially abrogated the hyperphagic obesity and hyperglycemia of *Lepr*-null (KO) mice (Fig. 5B-E; Fig. S8). This phenotype contrasts with the minimally impacted food intake and body weight following the reactivation of *Lepr* expression in *Pomc* neurons or in the entire ARC of otherwise *Lepr*-null mice (30, 31). While Re^Glp1r^ males displayed decreased blood glucose compared to KO mice, it remained elevated compared to controls (Fig. 5E).

**Fig. 5.**
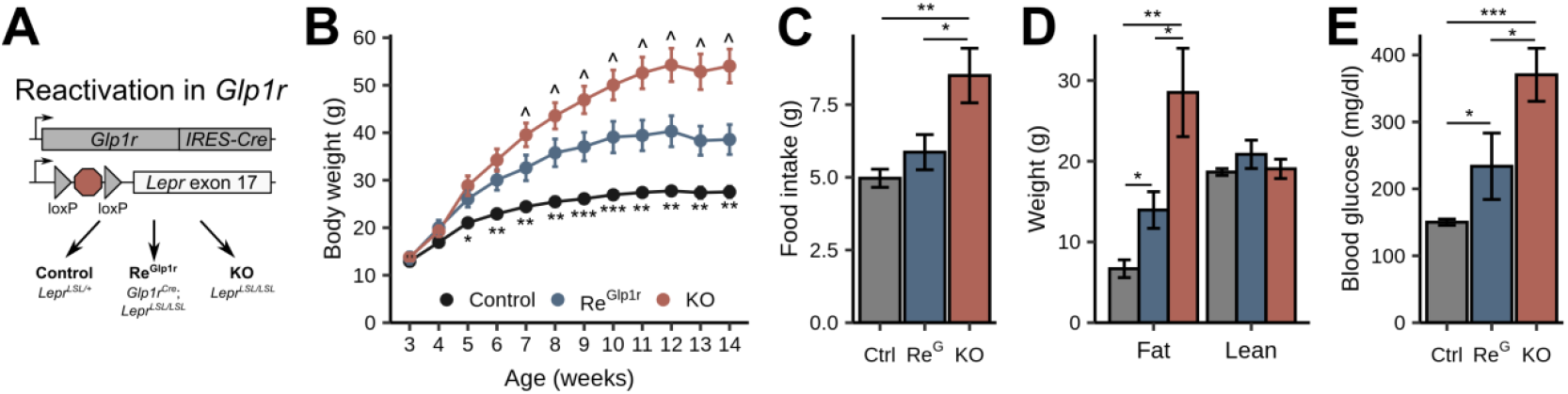
Leptin action via LepRb^Glp1r^ cells suppresses food intake and body weight. (**A**) Experimental diagram for generating Re^Glp1r^, KO, or control animals. (**B**) Body weight, (**C**) food intake, (**D**) body composition, and (**E**) blood glucose in Control (black, *n* = 8–12), KO (red, *n* = 5–11), and Re^Glp1r^ (blue, n = 5–8) male mice. For panel B, * symbol refers to Control vs. Re^Glp1r^ and ^ to KO vs. Re^Glp1r^. For all panels, 3 symbols: *P* < 0.001, 2 symbols: *P* < 0.01, 1 symbol: *P* < 0.05; all statistical significance determined by permutation test.

*Lepr* expression in the larger set of GABAergic (marked by the expression of the vesicular GABA transporter (vGAT; encoded by *Slc32a1*)) LepRb neurons plays a crucial role in the control of energy balance by leptin (10, 14). Sizeable GABAergic LepRb populations include LepRb^Glp1r^, LepRb^Agrp^, LepRb^Ghrh^, and LepRb^Nts^ cells (Fig. 6A); the early developmental ablation of *Lepr* from LepRb^Agrp^, LepRb^Ghrh^, or LepRb^Nts^ cells populations only modestly affects energy balance, however (9, 10, 21). We thus set out to determine whether LepRb^Glp1r^ cells might represent a GABAergic LepRb population crucial for the control of energy balance.

**Fig. 6.**
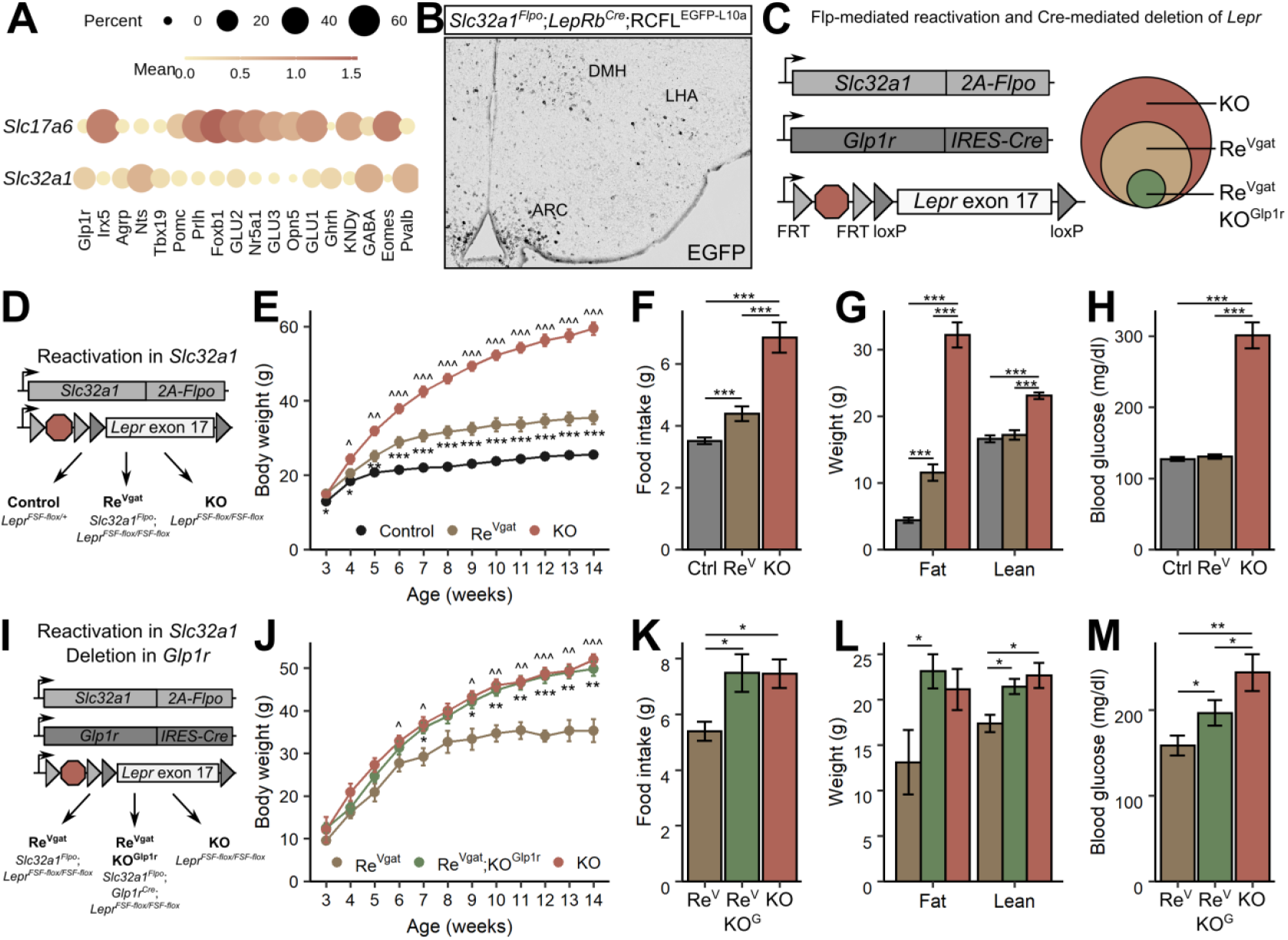
GABAergic Lepr^Glp1r^ cells mediate leptin action. (**A**) Expression of *Slc17a6* (vGlut2) or *Slc32a1* (vGat) for LepRb^Sun1-sfGFP^ populations. (**B**) Representative image of GFP-IR (black) in the hypothalamus of *Slc32a1*^*Flpo*^;*LepRb*^*Cre*^ mice on a Flp- and Cre-dependent GFP reporter background. (**C**) Strategy for testing the role of LepRb^Glp1r^ cells in leptin effects via GABAergic neurons. (**D**–**H**) Sufficiency of *Lepr* in *Slc32a1*-expressing cells for the control of energy balance. (**D**) Experimental diagram for generating *Slc32a1*^*Flpo*^;*Lepr*^*FSF-flox/FSF-flox*^ (Re^vGAT^), *Lepr*^*FSF-flox*^*/*^*FSF-flox*^ (KO), or Control animals. (**E**) Body weight, (**F**) food intake, (**G**) body composition, and (**H**) blood glucose in control (black, *n* = 18–22), KO (red, *n* = 12), and Re^vGAT^ (gold, *n* = 13–15) mice. For panel E, ^ symbol refers to Re^vGAT^ vs. KO, * to KO vs. Re^vGAT^. (**I**–**M**) Requirement for *Lepr* in GABAergic Glp1r cells for the control of energy balance. (**I**) Experimental diagram for generating Re^vGAT^, *Slc32a1*^*Flpo*^;*Glp1r*^*Cre*^;*Lepr*^*FSF-flox/FSF-flox*^ (Re^vGAT^KO^Glp1r^), or KO animals. (**J**) Body weight, (**K**) food intake, (**L**) body composition, and (**M**) blood glucose in Re^vGAT^ (gold, *n* = 4–7), KO (red, *n* = 10–20), and Re^vGAT^KO^Glp1r^ (green, n = 10–18) mice. For panel J, ^ symbol refers to KO vs. Re^vGAT^ and * refers to Re^vGAT^KO^Glp1r^ vs Re^vGAT^. For all panels E–M, 3 symbols: *P* < 0.001, 2 symbols: *P* < 0.01, 1 symbol: *P* < 0.05; all statistical significance determined by permutation test.

We generated a *Slc32a1*^*FlpO*^ mouse line to permit the selective targeting of GABAergic LepRb populations. Crossing *Slc32a1*^*FlpO*^ onto *Lepr*^*Cre*^ and a Cre- and Flp-dependent GFP reporter line promoted GFP expression in brain areas known to contain GABAergic, but not glutamatergic (e.g., VMH), LepRb neurons (Fig 6B). To examine the collective ability of GABAergic LepRb neurons to effect the control of energy balance by leptin in the absence of *Lepr* expression in other types of neurons, we crossed *Slc32a1*^*FlpO*^ onto the *Lepr*^*FSF-flox*^ mouse line (32), in which an FRT-flanked transcriptional STOP cassette lies upstream of *Lepr* exon 17, disrupting functional *Lepr* expression (Fig. 6C). Thus, the *Lepr*^*FSF-flox*^ allele is null for *Lepr* expression, but can be reactivated by Flp (32-34). Additionally, because *Lepr* exon 17 is floxed in *Lepr*^*FSF-Flox*^, Flp-reactivated *Lepr* expression from this allele can be deactivated by Cre (Fig. 6C). Global deletion of the FRT-flanked transcriptional blocker from this previously validated allele forms the basis for the widely-used *Lepr*^*flox*^ mouse line (see Figure 4)(8, 10, 17, 32). Hence, *Lepr* expression is restricted to GABAergic LepRb neurons in *Slc32a1*^*Flpo*^;*Lepr*^*FSF-flox/FSF-flox*^ (Re^Vgat^) mice; while *Lepr* expression is restricted to non-*Glp1r* GABAergic neurons in *Slc32a1*^*Flpo*^;*Glp1r*^*Cre*^;*Lepr*^*FSF-flox/FSF-flox*^ (Re^Vgat^KO^Glp1r^) mice (Fig. 6C).

Restoration of *Lepr* expression in GABAergic cells in Re^Vgat^ mice dramatically improved the body weight, food intake, and blood glucose phenotypes of *Lepr*-null (KO) mice (Fig. 6D–H). The remaining difference in energy balance between Re^Vgat^ and control mice did not result from insufficient Flp activity (Flp recombinase activity is notoriously lower than that of Cre), because *Slc32a1*^*Cre*^-mediated *Lepr* re-expression from the Cre-activated *Lepr*^*LSL*^ allele (30) displayed a similar phenotype (Fig. S9) (35). The difference in energy balance phenotype between Re^Vgat^ and control mice likely reflects the minor contribution of glutamatergic and other non-GABAergic LepRb cells to the control of energy balance (10, 14).

To determine the contribution of GABAergic LepRb^Glp1r^ cells to the rescue of energy balance control observed in Re^Vgat^ mice, we crossed *Glp1r*^*Cre*^ mice onto the Re^Vgat^ background to generate Re^Vgat^KO^Glp1r^ mice for comparison to Re^Vgat^ and KO mice (Fig 6I). Re^Vgat^KO^Glp1r^ mice phenocopied the hyperphagic obesity of KO mice, abrogating the restoration of leptin-mediated control of energy balance observed in Re^Vgat^ mice (Fig. 6J–M). While blood glucose in Re^Vgat^KO^Glp1r^ was elevated compared to Re^Vgat^ mice, it remained below that of KO mice, despite the equivalent body weight and adiposity of Re^Vgat^KO^Glp1r^ and KO mice. Thus, LepRb^Glp1r^ cells represent a key population of GABAergic LepRb neurons that are necessary and sufficient to mediate the control of food intake and energy balance by leptin, although they appear to play little role in the control of blood glucose and energy expenditure.

## Discussion

Our present findings define several previously unrecognized conserved populations of hypothalamic LepRb neurons, including LepRb^Glp1r^ neurons that play crucial roles in the control of the control of food intake and body weight by leptin. Additionally, LepRb^Glp1r^ neurons represent the previously-postulated population of DMH that plays critical roles in the control of energy balance. The important roles for these neurons in energy balance identify them as potential targets for obesity therapy.

Our data suggest that LepRb^Glp1r^ neurons predominantly control food intake; indeed, ablation of LepRb from these cells increased feeding and adiposity, while energy expenditure was appropriate for body weight in these mice. Furthermore, blood glucose remained normal in the KO^Glp1r^ mice despite their adiposity, and remained lower than KO mice in Re^Vgat^KO^Glp1r^ mice, despite their equivalent body weight and adiposity. We also speculate that KO^Glp1r^ mice might display normal endocrine function; indeed, we routinely breed KO^Glp1r^ mice (data not shown), which is not possible with KO mice (36).

While the downstream neural targets of LepRb^Glp1r^ neurons remain unknown, the specificity of these cells for the control of food intake during leptin action suggests different neural mechanisms of action for LepRb^Glp1r^ cells than for LepRb^POMC^ or LepRb^Agrp^ cells (the early developmental ablation of LepRb from which impacts the control of energy expenditure and/or glucose homeostasis more strongly than food intake). While LepRb^Glp1r^ cells might project to the ARC to modulate expressing POMC and/or AgRP neurons (including non-LepRb expressing subpopulations of these cells that may play distinct roles compared to LepRb^Pomc^ and LepRb^Agrp^ cells (37)), the specificity of LepRb^Glp1r^ neurons for food intake control mimics the function of *Mc4r*-expressing neurons of the paraventricular hypothalamic nucleus (PVH) (38), suggesting *Mc4r*-expressing or similarly acting neurons of the PVH as potential targets for LepRb^Glp1r^ cells.

Like ARC POMC and AgRP neurons, LepRb^Glp1r^ neurons presumably respond to stimuli other than leptin. Indeed, peripherally-administered semaglutide (a GLP1R agonist that drives 15% weight loss in a recent phase 3 clinical trial (39)) accumulates in the DMH (40), consistent with the potential modulation of LepRb^Glp1r^ neurons by exogenous GLP1R agonists and/or endogenous GLP-1. Hence, it will be important to determine the potential roles for LepRb^Glp1r^ cells for the Glp1r-dependent suppression of food intake.

Our snRNA-seq analysis also identified several additional novel populations of hypothalamic LepRb neurons, including LepRb^Tbx19^, LepRb^Foxb1^, and LepRb^Opn5^ cells-all of which express substantial *Lepr* and some of which exhibit clear cross-species conservation. In future studies it will be important to define the roles and mechanisms of action for these LepRb populations in leptin action. Similarly, the analysis of more LepRb neurons from LepRb^Sun1-sfGFP^ mice may reveal additional conserved populations from within the GABA and GLU1-3 populations that currently exhibit poor differentiation; it will be important to determine the roles for these populations as they are identified.

## METHODS

### Resource availability

Further information and requests for resources and reagents should be directed to and will be fulfilled by the lead contact, Martin G. Myers, Jr. All materials related to this study, including *Slc32a1*^*Flpo*^ mice produced for this study, are available upon reasonable request. All data will be publicly available upon publication. Sequencing data, count matrix files, and metadata are available through the Gene Expression Omnibus (GSE172463). Analysis scripts for all results presented in the paper, including mouse phenotyping data, are available at github.com/alanrupp/rupp-jclininvest-2022.

### Animals

Mice and rats were bred in the Unit for Laboratory Animal Medicine at the University of Michigan. Animals were provided with *ad libitum* access to food (Purina Lab Diet 5001) and water in temperature-controlled (25°C) rooms on a 12 h light-dark cycle with daily health status checks.

*Glp1r-IRES-Cre* mouse line (Jax: 029283) was a gift of Dr. Stephen Liberles (41). *Lepr*^*flox*^ and *Lepr*^*neo*^ lines were gifts of Dr. Streamson Chua (32). *Lepr*^*loxTB*^ mouse line (Jax: 018989) was a gift of Dr. Joel Elmquist (30). *ROSA26* ^*CAG-LSL-eGFP-L10a*^ mice and *Lepr*^*Cre*^ mice (Jax: 032457) have been described previously (42). *Slc32a1*^*Flpo*^ mice were generated for this study (see below for details).

#### Generation of *Slc32a1*^*Flpo*^ mouse line

Slc32a1-2A-Flpo (*Slc32a1*^*Flpo*^) mice were generated using recombineering techniques as previously described (43). Briefly, the Flpo transgene (AddGene plasmid #13793) and a loxP-flanked neomycin selection cassette were subcloned after 2A self-cleaving peptide. The 2A-Flpo-neomycin cassette was then targeted 3bp downstream of the stop codon of *Slc32a1* in a bacterial artificial chromosome. The final targeting construct containing the *Slc32a1*-*2A-Flpo* neomycin cassette and 4kb of flanking genomic sequence on both sides was electroporated into ES cells followed by neomycin selection. Appropriately targeted clones were identified by quantitative PCR and confirmed by southern blot analysis. Targeted clones were expanded and injected into blastocysts by the University of Michigan Transgenic Core. Chimeric offspring were then bred to confirm germline transmission of the *Slc32a1*-*2A-Flpo* allele; the neomycin selection cassette was removed by breeding to the E2A-Cre deleter strain (Jax: 003724).

#### Longitudinal study

Mice were maintained on standard chow diet and body weight was measured weekly from weaning. Blood glucose was measured biweekly from age 5 weeks. Some cohorts were single-housed to measure food intake weekly; other cohorts were single-housed at 12 weeks old and food intake was measured for 2 weeks. Mice were subjected to body composition measurements (Bruker Minispec LF 90II) at 16 weeks.

#### Tissue prep, cDNA amplification and library construction for 10x snRNA-seq

Mice and rats were euthanized using isoflurane and decapitated, the brain was subsequently removed from the skull and sectioned into 1mm thick coronal slices using a brain matrix. The mediobasal hypothalamus was dissected and flash frozen in liquid N_2_.

On the day of the experiment, frozen tissue (from 2–3 mice or 1 rat) was homogenized in Lysis Buffer (EZ Prep Nuclei Kit, Sigma) with Protector RNAase Inhibitor (Sigma) and filtered through a 30mm MACS strainer (Myltenti). Strained samples were centrifuged at 500 rcf x 5 minutes and pelleted nuclei were resuspended in washed with wash buffer (10mM Tris Buffer pH 8.0, 5mM KCl, 12.5mM MgCl_2,_ 1% BSA with RNAse inhibitor). Nuclei were strained again and recentrifuged at 500rcf x 5 minutes. Washed nuclei were resuspended in wash buffer with propidium iodide (Sigma) and stained nuclei underwent FACS sorting on a MoFlo Astrios Cell Sorter. For LepRb-Sun1 experiments, GFP+ and PI+ nuclei were collected; for all other experiments only PI+ nuclei were collected. Sorted nuclei were centrifuged at 100rcf x 6 minutes and resuspended in wash buffer to obtain a concentration of 750 – 1200 nuclei/uL. RT mix was added to target ∼10,000 nuclei recovered and loaded onto the 10x Chromium Controller chip. The Chromium Single Cell 3’ Library and Gel Bead Kit v3, Chromium Chip B Single Cell kit and Chromium i7 Multiplex Kit were used for subsequent RT, cDNA amplification and library preparation as instructed by the manufacturer. Libraries were sequenced on an Illumina NovaSeq 6000 (pair-ended with read lengths of 150 nt).

#### snRNA-seq data analysis

The FASTQ files were mapped to the appropriate genome (Ensembl GRCm38 or Rnor_6.0) using cellranger to generate count matrix files and data was analyzed in R. Macaque count matrix files were downloaded from GEO dataset GSE172203. Genes expressed in at least 5 cells in were retained. Cells with at least 600 detected genes were retained. Doublets were scored using Scrublet (44); clusters with a median doublet score > 0.3 or individual cells with scores >0.3 were removed.

The data was then normalized using scran (45) and centered and scaled for each dataset independently and genes that were called variable by Seurat *FindVariableFeatures* were input to PCA. The top PCs were retained at the “elbow” of the scree plot (normally 15-30, depending on the dataset) and then used for dimension reduction using UMAP and clustering using the Seurat *FindNeighbors* and *FindClusters* functions. *FindClusters* was optimized for maximizing cluster consistency by varying the resolution parameter from 0.2 upward in steps of 0.2 until a maximal mean silhouette score was found. Clusters were then hierarchically ordered based on their Euclidean distance in PC space and ordered based on their position in the tree.

Cell types were identified by projecting labels from a published hypothalamic single cell RNA-seq dataset (GSE87544) using the Seurat CCA method. Cells were labeled by the highest scoring cell type for the cell and its 14 nearest neighbors (from Seurat SNN). Neuron cluster names were chosen based on genes found in unbiased marker gene search (Seurat *FindMarkers*). Clusters without unique marker genes were labeled by their neurochemical identity (GABA or GLU).

#### TRAP-seq analysis

Published TRAP-seq data (GSE162603) was used to identify signatures of hypothalamic Lepr cells. Enriched genes were determined using DESeq2 including an effect of sample pair in the model to account for pairing of the bead–sup samples (∼ Pair + Cells). Effects of leptin were identified using only Bead samples with 10-hr treatment (PBS or Leptin) and a DESeq2 model of ∼ Genotype (WT or *ob*/*ob*) + Treatment.

Association of leptin-regulated genes and LepRb^Sun1-sfGFP^ populations was performed by projecting the scaled expression of significant leptin-regulated genes into principal component space. The association score is the magnitude of the first principal component.

#### Species integration

Mouse, rat, and macaque datasets were processed in the same way as for LepRb^Sun1-sfGFP^ including identifying celltypes and neuron clusters. For rat and macaque, the data was first converted into 1:1 orthologs using the mouse Ensembl annotation from biomaRt. To project LepRb^Sun1-sfGFP^ cells onto the UMAP embeddings of each species, we used Seurat’s *MapQuery* function with the CCA reduction. To identify ortholog populations, we generated a combined dataset all species and only their ortholog genes. Datasets were harmonized using Harmony and then clustered as previously. Clusters in which 80% of cells from each LepRb^Sun1-sfGFP^ population were labeled based on the LepRb^Sun1-sfGFP^ name. If more than one LepRb^Sun1-sfGFP^ population had 80% of its cells associated with the conserved cluster, these clusters were subclustered and the subcluster containing the most LepRb^Sun1-sfGFP^ cells from each cluster was assigned that name. Leptin-response scores were generated using the same approach as for the LepRb^Sun1-sfGFP^ dataset.

#### Immunostaining

Mice were euthanized with isofluorane and then perfused with phosphate buffered saline for five minutes followed by an additional five minutes of 10% formalin. For immunohistochemistry of phosopho-STAT3, mice were injected with leptin (5 mg/kg) 1 hour prior to euthanasia. Brains were then removed and post-fixed in 10% formalin overnight hours at room temperature, before being moved to 30% sucrose until sunk. Brains were then sectioned as 30 mm thick free-floating sections and stained. Sections were treated sequentially with 1% hydrogen peroxide/0.5% sodium hydroxide, 0.3% glycine, 0.03% sodium dodecyl sulfate, and blocking solution (PBS with 0.1% triton, 3% normal donkey serum; Fisher Scientific). Immunohistochemical and immunofluorescent staining. The sections were incubated overnight at room temperature in was preformed using standard procedures using the primary and secondary antibodies below. The following day, sections were washed and incubated with either biotinylated (1:200 followed by avidin-biotin complex (ABC) amplification and 3,3-diaminobenzidine (Thermo Scientific) or fluorescent secondary antibodies with species-specific Alexa Fluor-488 or -568 (Invitrogen A-11039 or A-11011, 1:250) to visualize proteins. Primary antibodies used include: GFP (1:1000, #1020, Aves Laboratories) and phosopho-STAT3 (1:500, #9145, Cell Signaling). Images were collected on an Olympus BX51 microscope. Images were background subtracted and enhanced by shrinking the range of brightness and contrast in ImageJ.

#### RNAscope

*In situ* hybridization was performed with the RNAscope® Multiplex Fluorescent Assay v2 from Advanced Cell Diagnostics (ACD) combined with Tyramide Signal Amplification technology (TSA™) and dyes from Akoya Biosciences. Fresh frozen brains collected from 38 10 week old Lepr^eGFP-L10a^ female mice were sliced on a cryostat at 16μm and mounted directly onto Superfrost Plus slides (Fisher). The tissue was fixed in 10% neutral buffered formalin for 15 min and then dehydrated in ethanol.

Endogenous peroxidase was blocked with H_2_O_2_ for 10 min, washed in DEPC water and then tissue was gently digested for 30 min at room temperature using Protease IV from the ACD kit. The tissue was incubated for 2 h at 40°C using probes targeting EGFP, Mm-Ebf1, Mm-Glp1r (#418851), and ACD’s 3-plex RNAscope® Positive Control Probes for mouse tissue. After hybridization, probes were labeled with Akoya dyes using TSA technology. Sections were counterstained with DAPI and cover slipped with ProLong Gold antifade mounting medium.

### Quantification and statistical analysis

All plotting and statistical analysis was performed using R 4.1.1. Specific statistical tests are listed in the figure legends.

### Study approval

All procedures performed were approved by the University of Michigan Committee on the Use and Care of Animals and in accordance with Association for the Assessment and Approval of Laboratory Animal Care and National Institutes of Health guidelines.

## Author contributions

Conceptualization, ACR and MGM; Methodology, ACR, AJT, AHA, and MGM; Formal Analysis, ACR; Investigation, ACR, AJT, AHA, CT, SRL, MAK, and AM; Resources, DPO; Data Curation, ACR; Writing – Original Draft, ACR and MGM; Writing – Review & Editing, all authors; Visualization, ACR; Supervision, DPO, CL, and LBK; Funding Acquisition, PK and MGM.

## Acknowledgments

We thank members of the Myers and Olson labs for helpful discussions, and Qing Zhu from the Molecular Genetics Core for mouse line production. We thank Steve Liberles (Harvard Medical School) for sharing *Glp1r*^*cre*^, and Joel Elmquist (University of Texas Southwestern) and Streamson Chua (Albert Einstein College of Medicine) for sharing the *Lepr*^*LSL*^ and *Lepr*^*FSF-flox*^ alleles. We thank Carey Lumeng and members of his lab for insights into sorting nuclei for snRNA-seq.

## Funding

Research support was provided by NIH DK056731 (MGM) and from the Michigan Diabetes Research Center (NIH P30 DK020572, including the Molecular Genetics and Animal Studies Cores), the Marilyn H. Vincent Foundation (MGM), and Novo Nordisk (to MGM).

## Data and materials availability

All data will be publicly available upon publication. Sequencing data, count matrix files, and metadata are available through the Gene Expression Omnibus (GSE172463). Analysis scripts and data for all results presented in the paper, including mouse phenotyping data, are available at github.com/alanrupp/rupp-jclininvest-2021. Mouse models will be made available upon reasonable request. Reagents will be made freely available to academic laboratories.

## SUPPLEMENTAL FIGURES

**Figure S1:**
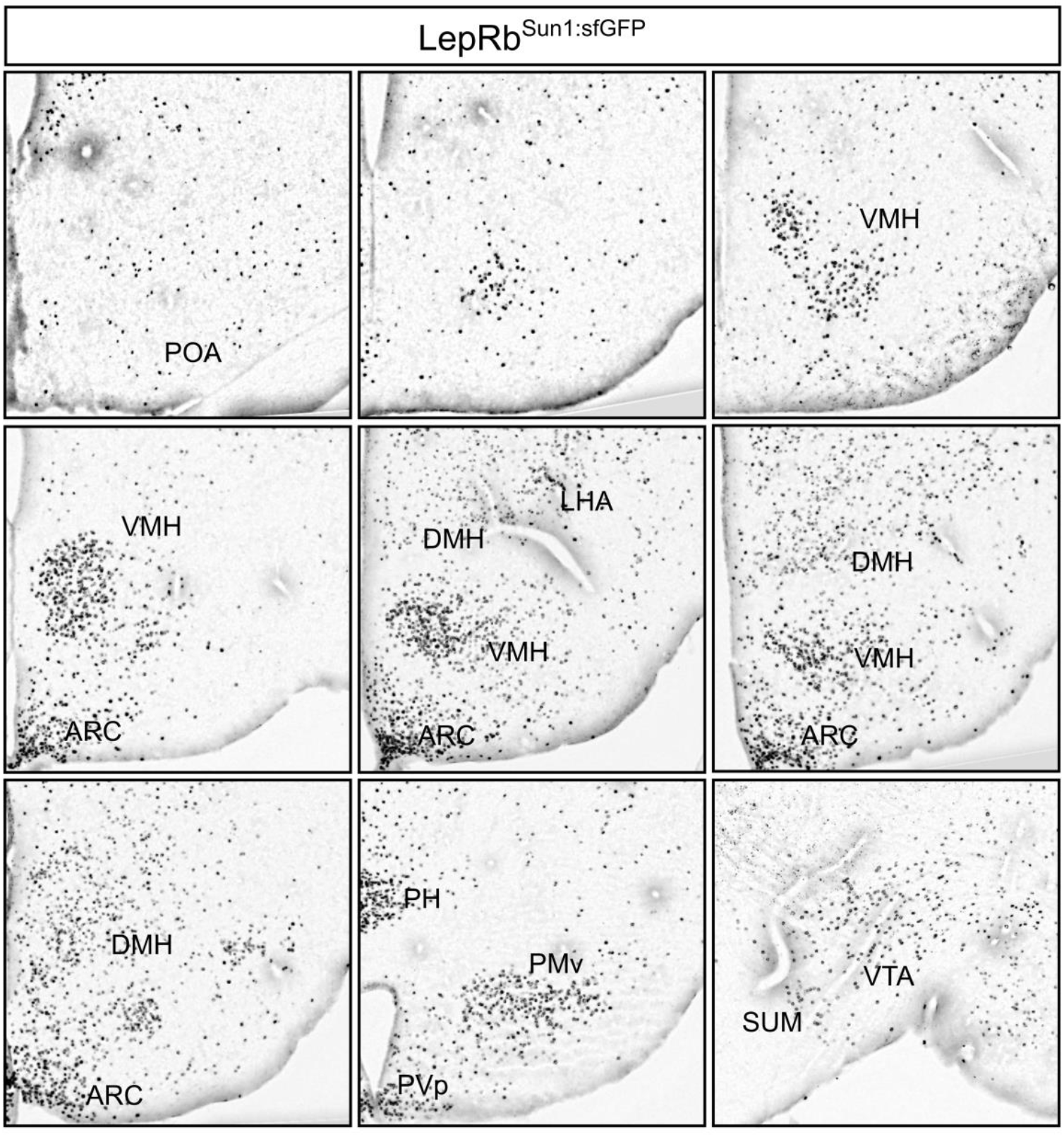
Sun1-sfGFP expression in the hypothalamus of LepRb^Sun1-sfGFP^ mice. Representative images of GFP-IR (black) in LepRb^Sun1-sfGFP^ hypothalamus, from rostral to caudal going left to right and then down rows.POA, preoptic area; LHA, lateral hypothalamic nucleus; ARC, arcuate nucleus, PH, posterior hypothalamus; PMv, ventral premammilary nucleus; PVp, posterior periventricular hypothalamic nucleus; SUM, supramammillary nucleus; VTA, ventral tegmental area.

**Figure S2:**
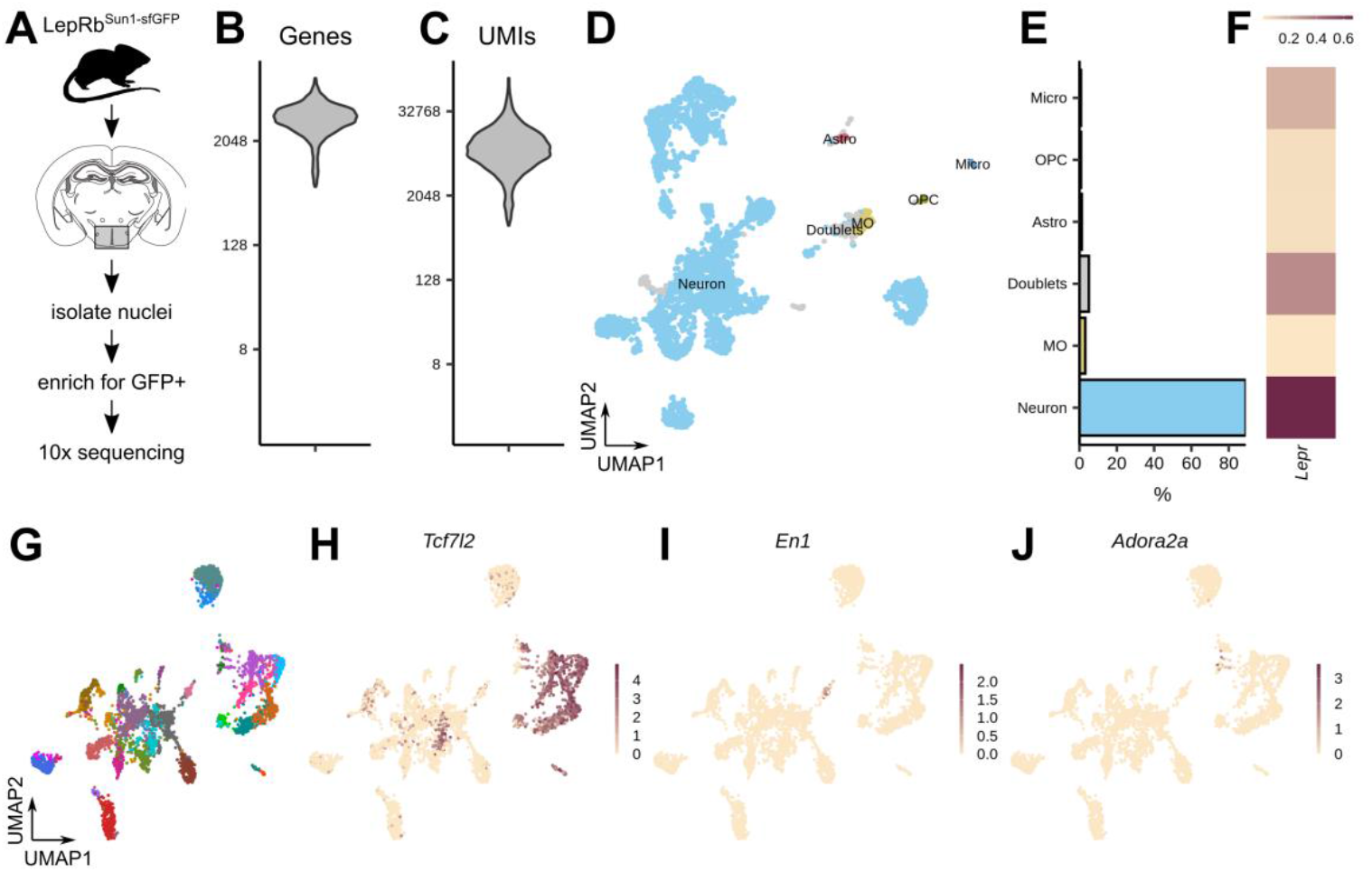
Classification of nuclei from LepRb^Sun1-sfGFP^ dataset. **(A)** Experimental design. **(B-C)** Distribution of **(B)** genes and **(C)** UMIs detected per cell. **(D)** UMAP projection of all cells colored by cell type. (E) Celltype membership breakdown. (F) Mean Lepr expression in each cell type. (G) UMAP projection of all neurons, colored by cluster. **(H-J)** Expression of extrahypothalamic markers **(H)** *Tcf7l2*, (I) *En1*, and **(J)** *Adora2a* across all neurons.

**Figure S3:**
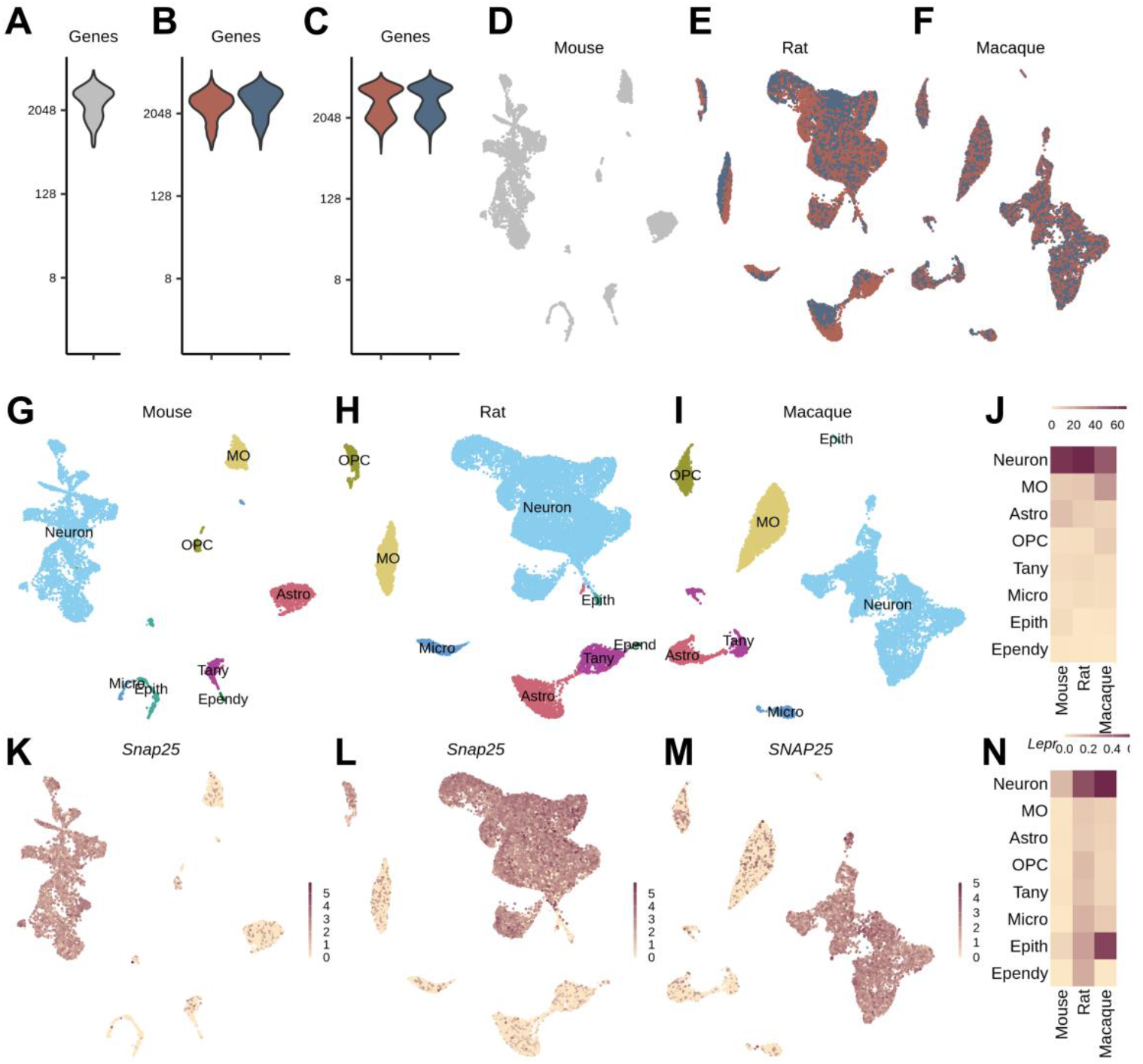
Classification of nuclei from mouse, rat, and macaque datasets. **(A-C)** Distribution of genes per cell for the samples from **(A)** mouse, **(B)** rat, and **(C)** macaque. **(D-F)** UMAP projection of all cells from **(D)** mouse, **(E)** rat, and **(F)** macaque, colored by sample. **(G-l)** UMAP projection of all cells from **(G)** mouse, **(H)** rat, and **(I)** macaque, colored by cell type. **(J)** Celltype membership breakdown. **(K-M)** Neuron marker *Snap25ISNAP25* expression across all cells in **(K)** mouse, **(L)** rat, and **(M)** macaque. **(N)** Mean *LeprILEPR* expression in each cell type.

**Figure S4:**
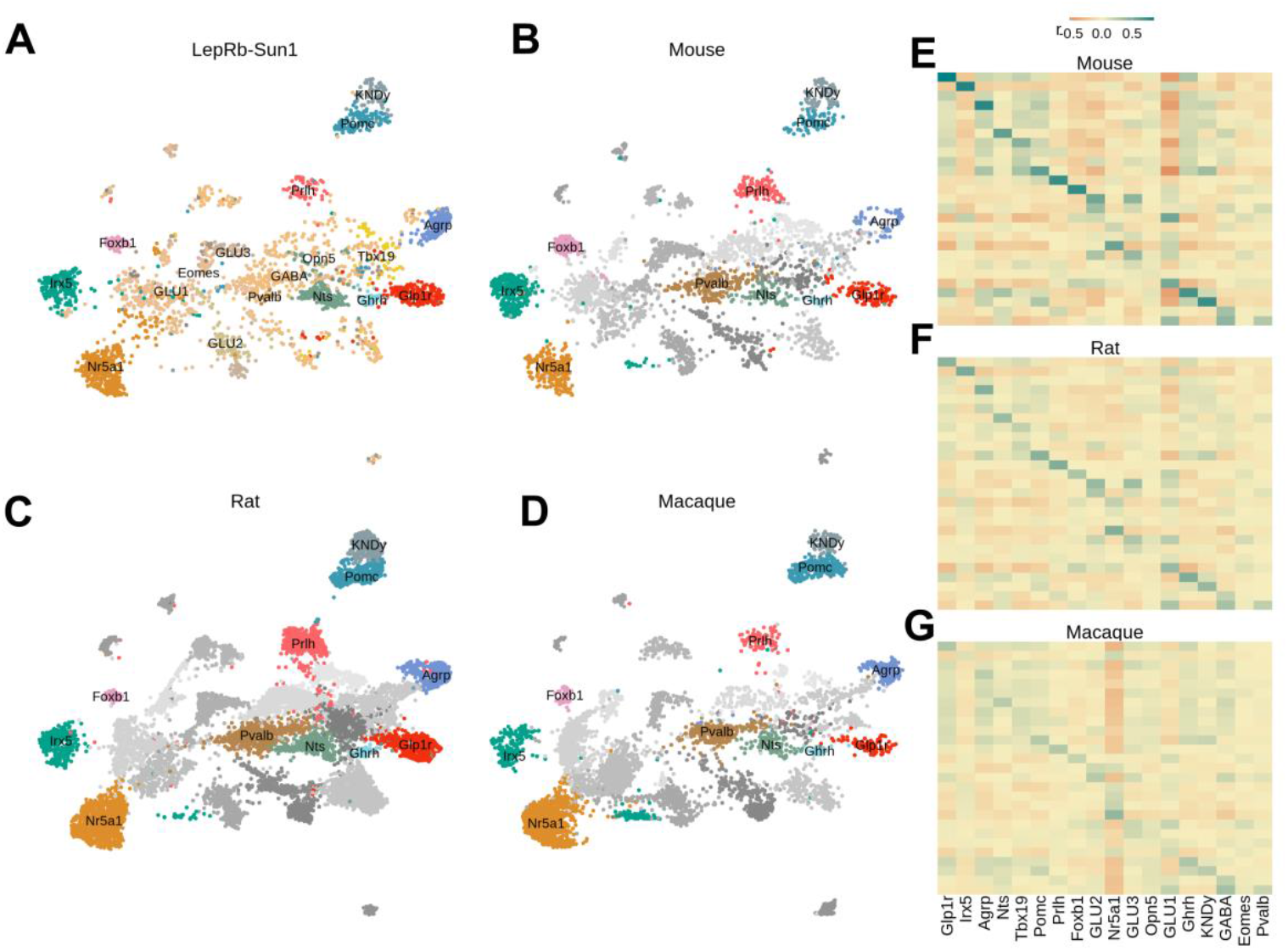
Conservation of LepRb populations across species. **(A-D)** UMAP projection of harmonized dataset of neurons from all species, colored by harmonized cluster. **(E-G)** Pairwise expression correlation of all marker genes in all LepRb^Sun1-sfGFP^ populations (x-axis) and harmonized populations (y-axis) for each species.

**Figure S5:**
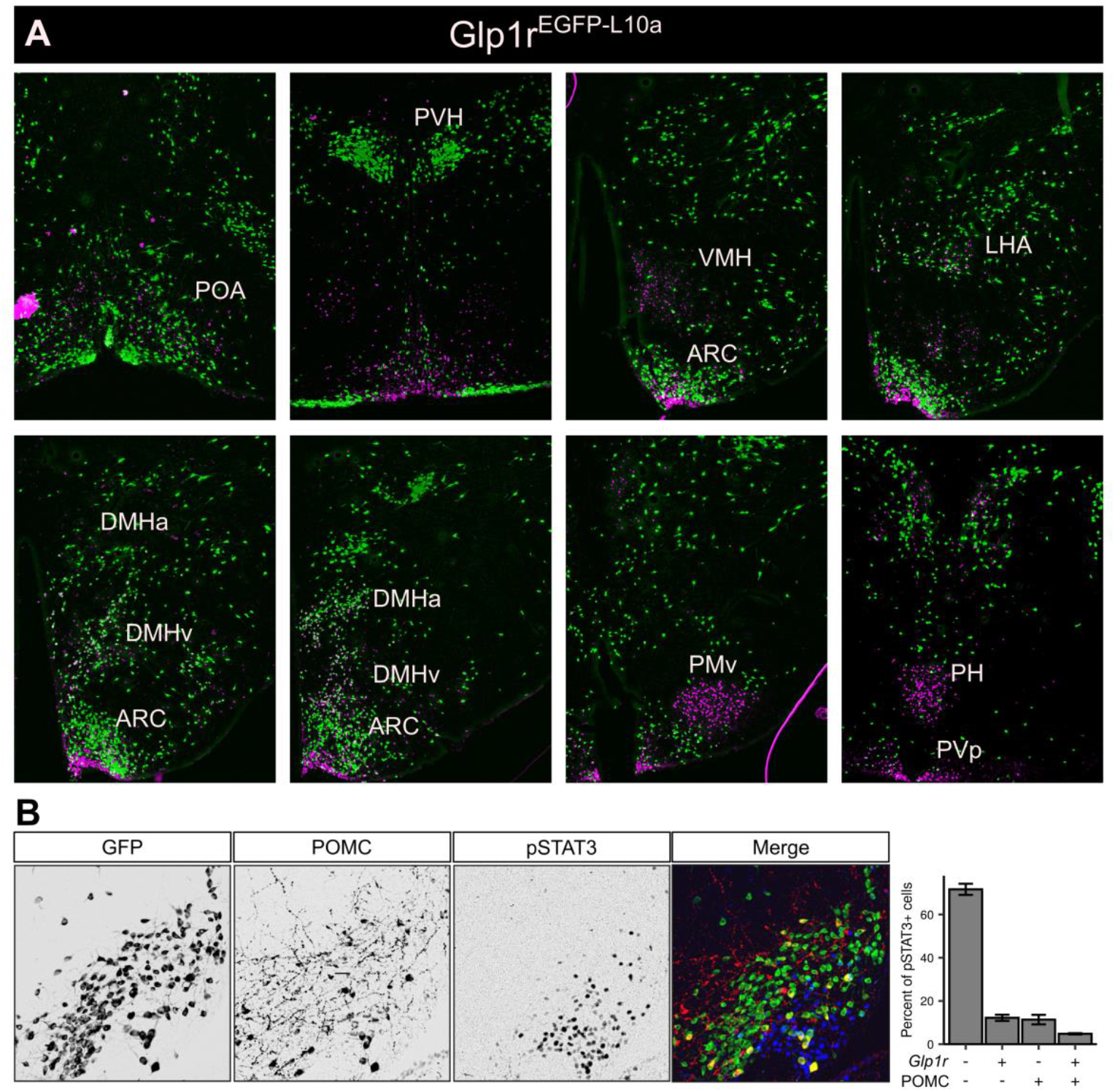
Colocalization of *Glplrand Lepr in* the DMH and lack of colocalization of LepRb^Glp1r^ neurons with POMC. **(A)** Representative images showing GFP-IR (green) and leptin-stimulated pSTAT3-IR (a marker for LepRb, magenta) throughout the hypothalamus of a Glp1 r^EGFP-L10a^ mouse. **(B)** Representative image showing GFP-IR (left; green in merged panel), POMC-IR (second from left, red in merged panel), pSTAT3-IR (third panel from left, blue in merged panel), and merged images from the ARC of a Glp 1 r^EGFP-L10a^ mouse. Right: Graph showing quantification of the percent of ARC LepRb cells that colocalize with POMC and/or GFP-IR *(Glp1r)*. Mean ± SEM is shown, n = 3.

**Figure S6:**
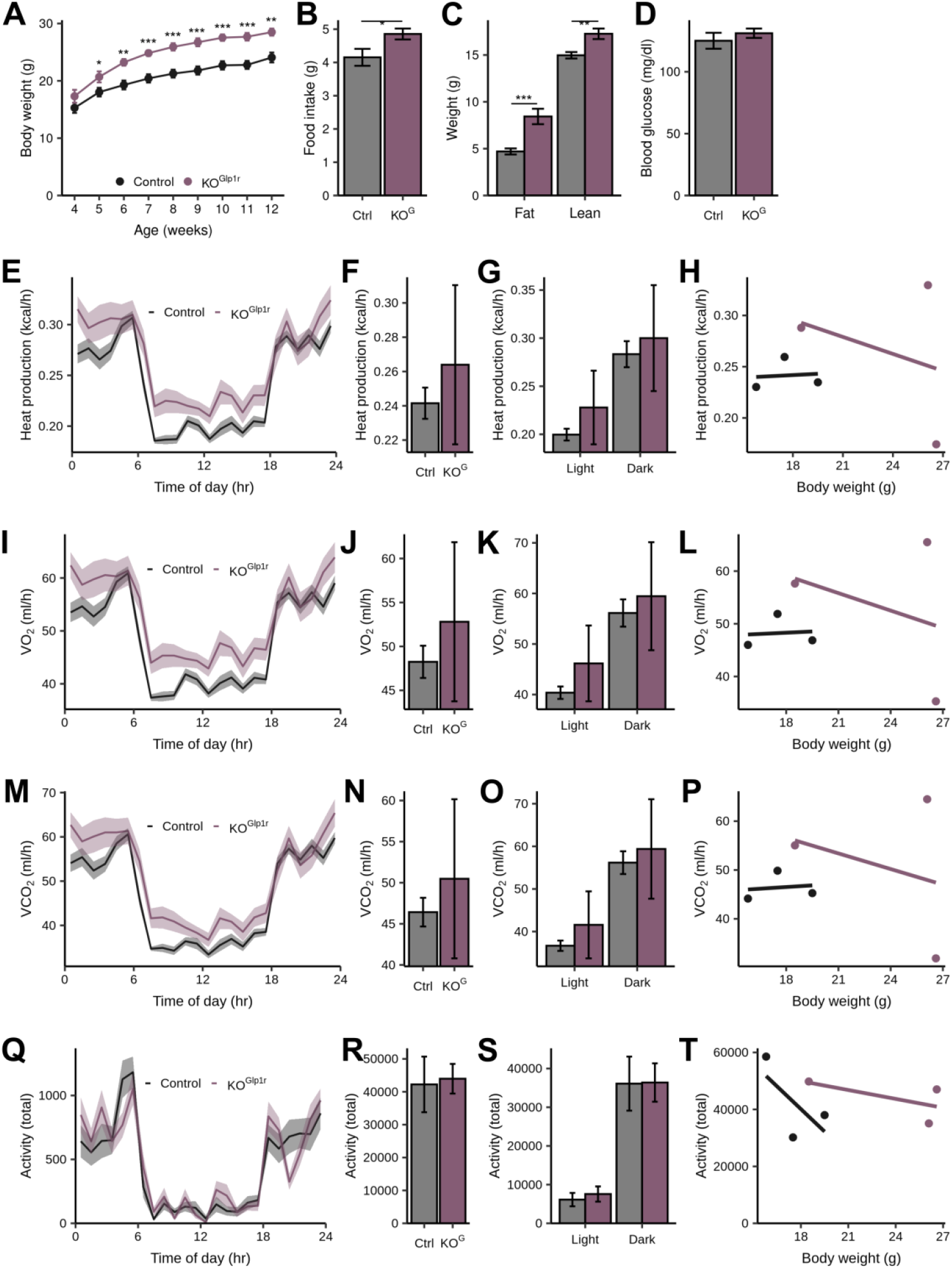
Metabolic parameters of female KO^Gp1r^ mice. **(A-D) (A)** Body weight, **(B)** food intake, and **(C)** fat mass, but not **(D)** ad libitum blood glucose, are all elevated in KO^Glp1r^ female mice *(Glp1r*^*Cre*^*;Lepi*^*flox/flox*^, *n =* 7−9) relative to littermate controls (Lep^*flox/flox*^, *n* = 10-14). **(E-T)** Energy expenditure is not reduced in KO^Glp1r^ female mice (*n* = 3) relative to littermate controls (*n* = 3). Individual parameters are in rows, with the columns from left to right corresponding to data across the day, binned by 1 hour, daily average, daily average split by light/dark cycle, and daily average correlation with body weight. All data represents mean ± SEM, *** P < 0.001, ** P < 0.01, * P < 0.05 by permutation test.

**Figure S7:**
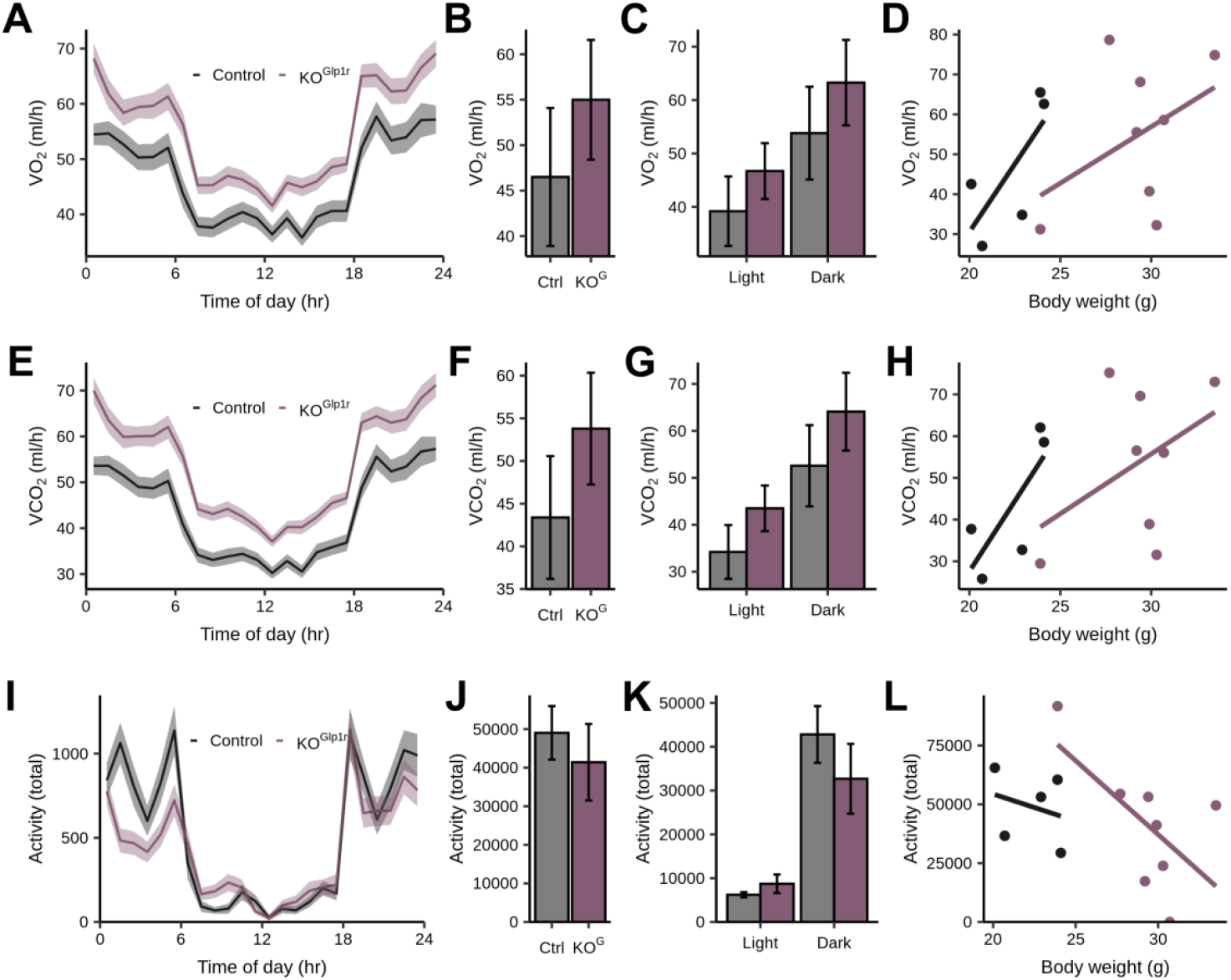
Energy expenditure parameters of male KO^Glp1r^ mice. Energy expenditure is not reduced in KO^Glp1r^ male mice *(n =* 8) relative to littermate controls *(n =* 5). Individual parameters are in rows, with the columns from left to right corresponding to data across the day, binned by 1 hour, daily average, daily average split by light/dark cycle, and daily average correlation with body weight. All data represents mean ± SEM, *** P < 0.001, ** P < 0.01, * P < 0.05 by permutation test.

**Figure S8:**
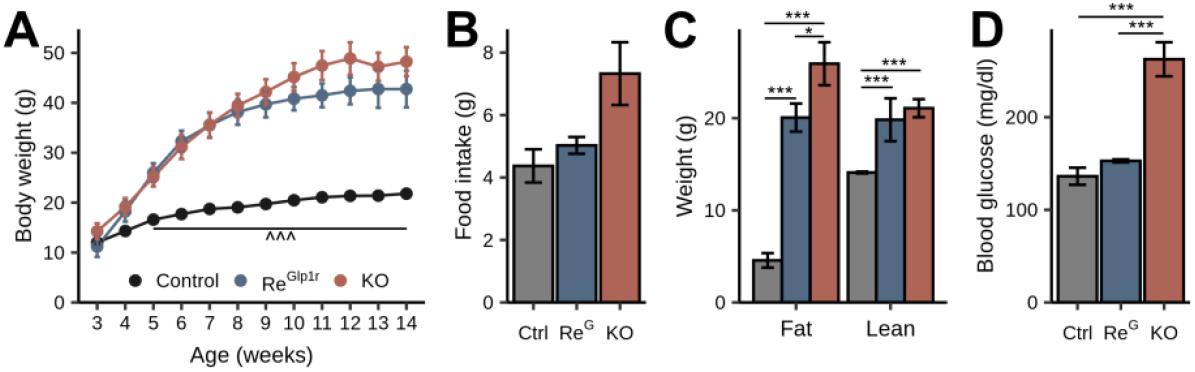
Metabolic parameters of female Re^Glp1r^ mice. **(A-D) (A)** Body weight, **(B)** food intake, **(C)** body composition, and **(D)** *ad libitum* blood glucose in Re^Glp1r^ female mice (blue, *Gip1r*^*Cre*^ *Lepr*^*LSL/LSL*^, n = 3−4) compared to llittermate controls (black, Lepr^*LSL/LSL*^) and LepRb null animals (red, KO, *Lepr*^*LSL/LSL*^) For longitudinal plot, *^^^ P <* 0.001 for Control vs. Re^Glp1r^. For **B-D**, *** P < 0.001, **P < 0.01, * P < 0.05 for comparisons labeled by the line. Data represents mean ± SEM.

**Figure S9:**
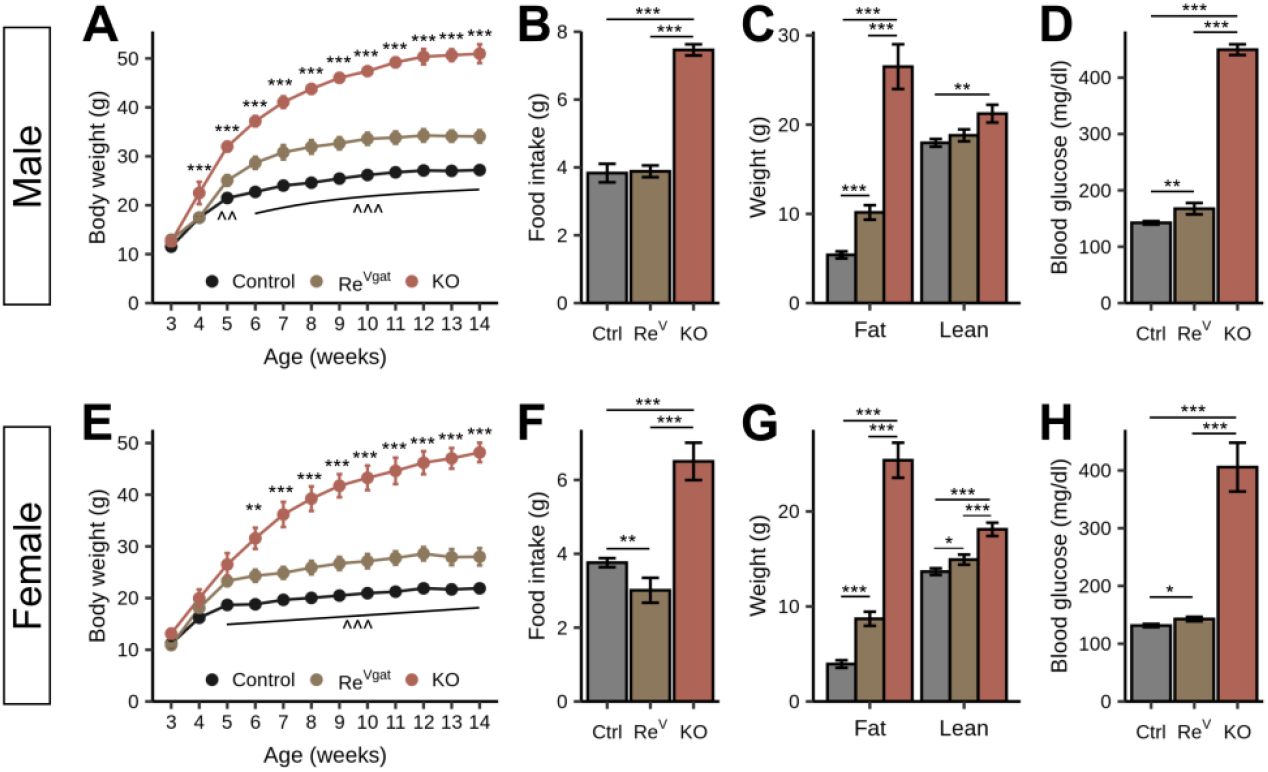
Cre-dependent rescue of Lepr in Slc32a1 cells displays similar phenotype as Flp-dependent rescue. **(A-D)** Male and **(E-H)** female body weight, food intake, body composition, and *ad libitum* blood glucose in Re^Vgat^ mice (gold, *Slc32a1*^*Cre*^*-,Lepr*^*LSL/LSL*^, *n = 7* males and 4 females) compared to littermate controls (black, *Lepr*^*LSL/+*^, *n* = 10 males and 8 females) and LepRb null animals (red, KO, *Lepr*^*LSL/LSL*^, n = 3 males and 6 females). For panels **A** and **E**, ^**^^^**^ *P* < 0.001, ^^^^ *P <* 0.01, ^^^ *P* < 0.05 for Control vs. Re^Vgat^ and *** *P <* 0.001, *\*\* P <* 0.01 for Re^Vgat^ vs. KO by permutation test. For panels **B, C, D, F, G**, and **H**, *** *P <* 0.001, ** *P <* 0.01, *\* P <* 0.05 by permutation test for comparisons labeled by the line. Solid data represents mean ± SEM.

## Notes

**Conflict of Interest Statement:** CL and LBK are employees of Novo Nordisk. DPO, PK, and MGM receive research funding from Novo Nordisk. MGM receives research funding from AstraZenica. The authors declare that no other conflicts exist.

### Competing Interest Statement

CL and LBK are employees of Novo Nordisk. DPO, PK, and MGM receive research funding from Novo Nordisk. MGM receives research funding from AstraZenica. The authors declare that no other conflicts exist.

